# *Aedes aegypti* CCEae3A carboxylase expression confers carbamate, organophosphate and limited pyrethroid resistance in a model transgenic mosquito

**DOI:** 10.1101/2023.08.16.553486

**Authors:** Beth C. Poulton, Fraser Colman, Amalia Anthousi, David B. Sattelle, Gareth J. Lycett

## Abstract

Insecticide resistance is a serious threat to our ability to control mosquito vectors which transmit pathogens including malaria parasites and arboviruses. Understanding the underlying mechanisms is an essential first step in tackling the challenges presented by resistance. This study aimed to functionally characterise the carboxylesterase, CCEae3A, the elevated expression of which has been implicated in temephos resistance in *Aedes aegypti* and *Aedes albopictus* larvae. Using our GAL4/UAS expression system, already established in insecticide-sensitive *Anopheles gambiae* mosquitoes, we produced transgenic *An. gambiae* mosquitoes that express an *Ae. aegypti* CCEae3A ubiquitously. This new transgenic line permits examination of CCEae3A expression in a background which does not express the gene and allows comparison with existing *An. gambiae* GAL4-UAS lines. Insecticide resistance profiling of these transgenic *An. gambiae* larvae indicated significant increases in resistance ratio for three organophosphate insecticides, temephos (5.98), chloropyriphos (6.64) and fenthion (3.18) when compared to the parental strain. Cross resistance to adulticides from three major insecticide classes: organophosphates (malathion, fenitrothion and pirimiphos methyl), carbamates (bendiocarb and propoxur) and pyrethroid (alpha-cypermethrin) was also detected. Resistance to certain organophosphates and carbamates validates conclusions drawn from previous expression and phenotypic data. However, detection of resistance to pirimiphos methyl and alphacypermethrin has not previously been formally associated with CCEae3A, despite occurring in *Ae. aegypti* strains where this gene was upregulated. Our findings highlight the importance of characterising individual resistance mechanisms, thereby ensuring accurate information is used to guide future vector control strategies.

**Author Summary:** Insecticides are vital disease control tools against pathogen-transmitting mosquitoes. However, they are becoming less effective as mosquitoes develop resistance. Among the molecular changes that contribute to resistance, increased production of enzymes that break down/sequester the insecticide is common. In *Ae. aegypti* mosquitoes, which spread many arboviruses, over-expression of the carboxylesterase enzyme, CEae3A, has been associated with resistance to certain insecticides used for vector control, particularly organophosphate compounds. However, multiple resistance enzymes/mechanisms are likely to be present in resistant mosquitoes at the same time. To examine the effect of CCEae3A expression in isolation, we utilised the *An. gambiae* mosquito with its convenient access to GAL4/UAS technology to regulate gene expression. This enabled production of CCEae3A in a normally insecticide-sensitive mosquito strain, permitting expression without interference from other resistance mechanisms. As anticipated, resistance to organophosphates was observed in larvae expressing CCEae3A. In adults, resistance was also found against compounds from organophosphate, carbamate and pyrethroid insecticide classes, including two compounds for which there had been no previous association. As well directly linking CCEae3A expression to specific insecticide resistance, this transgenic line can be included in a panel expressing alternative enzymes to screen new insecticidal compounds for liability to existing resistance mechanisms.

## 1 Introduction

Insecticide resistance in mosquito populations threatens to seriously undermine the successful interventions that have limited pathogen transmission and disease prevalence (1). Insecticides cause mortality through binding to vital protein targets, often neuronal, altering their activity to such a degree that death ensues. Resistance develops through inheritance of traits that reduce binding efficiency and/or the number of toxic molecules which access the target site (2). Such traits involve multiple molecular mechanisms including 1) amino acid changes in target sites that modify insecticide binding, 2) sequestration of insecticide away from target sites through binding with specific proteins, 3) cuticular barriers to insecticide uptake and 4) increased insecticide turnover by “metabolic’ enzymes resulting in products with reduced binding/toxicity and increased excretion.

In mosquitoes there are three large families of metabolic enzymes that have been strongly associated with detoxification of different chemical classes of insecticides; namely cytochrome P450s, glutathione-s-transferases (GST) and carboxylesterases (COE) (3). To identify specific family members that confer metabolic resistance, comparative high throughput transcriptomic and genomic approaches in resistant and sensitive strains are used to delineate potential candidates (4-9). Further evidence is then provided through demonstration of recombinant enzyme activity against insecticidal substrates. Finally, *in vivo* validation is sought through heterologous expression of candidates in *Drosophila melanogaster*, or more recently transgenic *An. gambiae* mosquitoes, and subsequent characterisation of insecticide resistance phenotypes (10-13). This approach permits identification, validation and understanding of the molecular drivers of resistance and provides crucial information on how best to curb the spread of these inheritable characteristics. It also provides mosquito lines with stable resistance phenotypes that can be used in screens to identify new insecticidal compounds that may circumvent established resistance mechanisms, thereby maintaining vector control. The development of *Aedes aegypti* as a model organism is underway (14) in the meantime we have adopted *An. gambiae* for the present work, where the UAS/GAL4 system is established and has been used to explore resistance genes including P450s and glutathione S transferase.

We have previously adapted the widely used GAL4-UAS to perform functional analysis of the most promising cytochrome mosquito P450 and GST candidates using the ubiGAL4/UAS system in *An. gambiae* (15). Overexpression of the enzymes, cyp6M2 or cyp6P3 with a near-ubiquitous tissue expression profile gave rise to resistance to pyrethroids and in the case of cyp6P3 also showed limited resistance to carbamates. P450 overexpression also increased sensitivity to a pro-insecticide OP, malathion, probably through metabolic activation. Meanwhile, GSTe2 transgenic analysis indicated a prominent role in DDT resistance as well as high levels of OP resistance (15). Extension of this work also helped to define a role for sensory appendage protein, Sap2, in pyrethroid resistance (16). Further work has shown the synergistic interaction of GSTe2 overexpression with the directed mutation of the pyrethroid target site, the voltage gated sodium channel (17).

This panel of metabolic gene overexpressing lines have been further used to screen a pipeline of new insecticides (and older agricultural insecticides that have recently been re-purposed for vector control) for their vulnerability to existing metabolic resistance mechanisms (18). To broaden the application of this technology, we wished to address some of the knowledge gaps in the analysis. These included the examination of candidates belonging to the unexplored COE family and to demonstrate that the *An. gambiae* ubi-GAL4/UAS system *An. gambiae* could be used to examine resistance in larval stages, since previous work had solely targeted adults.

Insecticidal aquatic larval control has primarily focused on Aedes mosquitoes, since the adults are more difficult to target because of their diurnal feeding habit. The OP temephos is the most commonly used chemical larvicide (19) due to its low mammalian toxicity and approved use in potable water sources. However, resistance has been observed worldwide. Resistance to OP insecticides in *An. gambiae* is strongly associated with the widespread insensitive ACE1 mutation G280S (previously denoted G119S) (20) but this modification is rarely observed in *Ae. aegypti* and *Ae. albopictus* mosquitoes as the specific codon used at that site requires two nucleotide changes rather than the single change needed in most insects (21).

One of the most common genes associated with OP resistance, and particularly resistance to temephos in Aedes mosquitoes, has been the COE encoding CCEae3A gene. 60-fold upregulation of CCEae3A (AAEL023844) was identified in temephos resistant *Ae. aegypti* from Thailand (22) and the French West Indies (23). Similarly, *Ae. albopictus* larvae selected with temephos in the laboratory displayed 27-fold CCEae3A upregulation in resistant mosquitoes (24). Amplification of CCEae3A copy number, presumably related to increased transcription, has also been associated with resistant phenotypes in both Aedes species (25). Moreover, recombinant CCEae3A from both Aedes species have been shown to sequester the OP and to metabolise the toxic oxon form of temephos to its less toxic product, [(4-hydroxyphenyl) sulfanyl] phenyl O,O-dimethylphosphorothioate (26).

As well as being linked to temephos resistance, CCEae3A upregulation has also been found in mosquitoes displaying resistance to permethrin, malathion, bendiocarb, fenitrothion and propoxur (22-25, 27-29). However, other target site and metabolic resistance mechanisms were also detected in these mosquitoes, which were found to be resistant to several insecticides.

To delineate and quantify the role of CCEae3A in larvae, and to allow comparison with similar experiments with P450 and GST enzymes in adults, we have used our established ubi-GAL4/UAS-target gene system to overexpress CCEae3A in transgenic *An. gambiae* mosquitoes. By this means, we determined whether overexpression of CCEae3A gives rise to resistance to different members of the OP insecticide class in larval stages, as well as other classes of public health adulticides.

## 2 Methods

### 2.1 GAL4/UAS and UAS:CCEae3A donor plasmid construction

We have previously developed a GAL4/UAS system in *An. gambiae* to express transgenes in a near ubiquitous expression pattern. Briefly, the system comprises a GAL4 driver line whose expression is controlled by the *An. gambiae* polyubiquitin promoter. The Ubi-GAL4 sequences are linked to a 3xp3 controlled eCFP to mark transgenics. This expression cassette is flanked by inverted PhiC31 AttP repeats and has been integrated into the genome by piggyBac transformation (14) (Fig 1A). The Ubi-GAL4 line acts both as a driver line, so that when crossed with responder lines carrying Gal4 responsive, UAS-geneX sequences, geneX is expressed in a ubiquitous pattern in the progeny, as well as a docking line carrying the inverted PhiC31 repeats. The docking aspect can be used to swap out the Ubi-GAL4-3xP3eCFP sequences with a new differentially marked expression cassette by double recombination at the AttP sites driven by PhiC31 integrase. In this instance, we have exchanged a cassette carrying a UAS:CCEae3A marked with eYFP (Fig 1B) to create a UAS responder line (expressing eYFP) (Fig 1C). If only a single recombination occurs between the donor plasmid and docking site, the whole of the donor plasmid integrates into that site and so in this instance, both the driver (eCFP) and responder (eYFP) cassettes are present on the same locus (Fig 1D).

**Fig 1.**
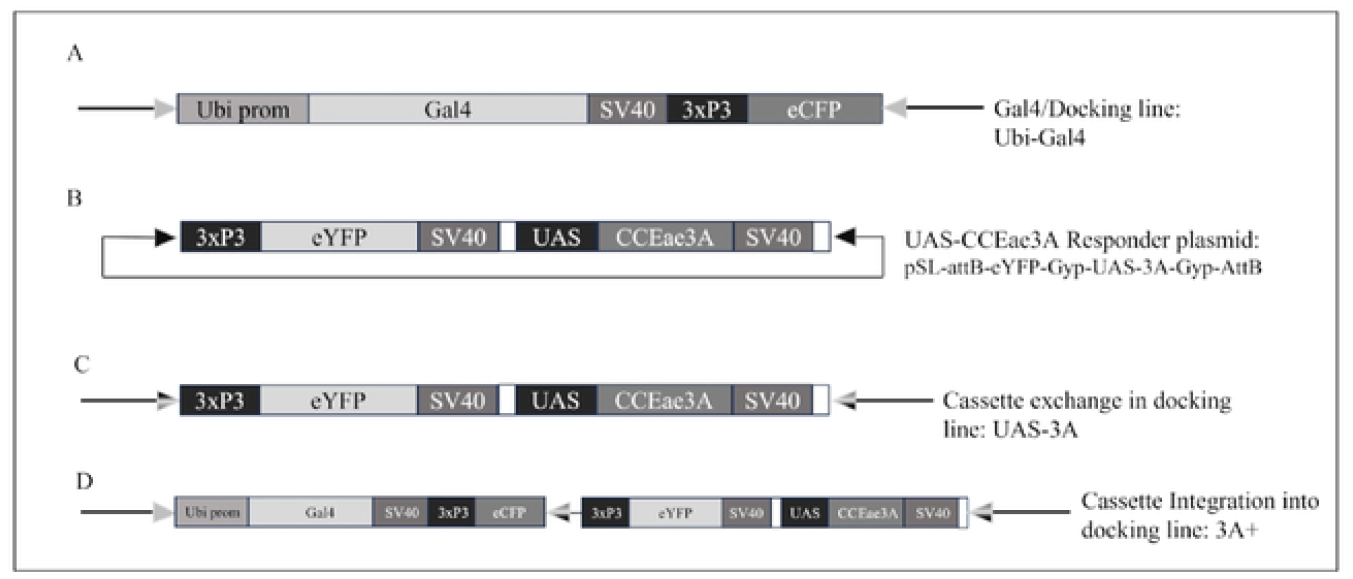
GAL4-UAS docking lines and transformation constructs. Representation of the (A) Ubi GAL4 docking line genome insertion, (B) UAS-CCEae3A responder plasmid, (C) UAS-3A cassette exchange genome insertion, (D) 3A+ cassette integration genome insertion. To achieve expression from the UAS-3A line (C), these are crossed with the UbiGal4 line, and CCEae3A is then expressed in the progeny. Since these insertions are in the same genome locus, only single copies of each transgene can be achieved when both are present. In D, both driver and responder transgenes are present on single allele, and the CCEae3A is thus transcribed ubiquitously and can be made homozygous by intercrossing. Pale arrowheads indicate AttP sites, dark arrowheads indicate AttB sites and hatched arrowheads indicate hybrid AttB/AttP sites.

To generate the UAS:CCEae3A responder plasmid (Fig1B), the CCEae3A cDNA (AAEL023844-RA) was first subcloned into a UAS plasmid (pSL*attB:YFP:Gyp:UAS14i:Gyp:attB) (10) previously described in (13). The 1669-bp CCEae3A cDNA sequence was amplified from a template derived from the temephos resistant Nakhon Sawan 2 *Ae. aegypti* strain (22, 26) - using primers CCEfor2 (GACTGGAATTCCATTATGTCCACTTTGGA) and CCErev2 (GTATTCTCGAGTCATTGCAATGCTCGATG). The amplified fragment and UAS plasmid (pSL*attB:YFP:Gyp:UAS14i:Gyp:attB) (10) were ligated following EcoRI and XhoI (NEB) digestion and purification. Positive clones were identified by colony PCR using CCEseqfor (ATTGTGGTGACGTTCAACTATCG) and CCEseqrev (GTATGTCAATTCATCCGCATGAGC) primers. Verification of the selected clone, pSL-attB-YFP-Gyp-UAS-3A-Gyp-attB, was confirmed by Sanger sequencing using the following primers: UASp (GCAAGGGTCGAGTCGAGCGGAGACTCTAGC), CCErev (CTCGAGTCATTGCAATGCTCGATG), CCEseqfor (ATTGTGGTGACGTTCAACTATCG) and CCEseqrev (GTATGTCAATTCATCCGCATGAGC).

### 2.2 Creation of *An. gambiae* lines by ΦC31-mediated cassette exchange

To generate the transgenic lines, a mixture of φC31 integrase-encoding helper plasmid (pKC40, 150 ng/μL) (30, 31) and 350 ng/μL UAS-CCEae3A responder plasmid were co-injected into embryos of the *An. gambiae* docking line Ubi-GAL4 using standard procedures (31). Emerged F0 larvae expressing eYFP fluorescence in their posterior tissues were selected by fluorescent stereomicroscopy (Leica FLIII) using a narrowband YFP filter (Leica). Virgin F0 individuals were outcrossed to 3-5 times the number opposite sex wild-type *An. gambiae* G3 in founder cages and females blood fed 4-5 days later. F1 larvae expressing eYFP (cassette exchange Fig1C) or eYFP + eCFP (cassette integration Fig1D) in nerve cord and eye were identified by fluorescence microscopy using YFP and CFP filters (LeicA), then allowed to grow to adulthood, before crossing with excess numbers of opposite sex Ubi-GAL4 or G3 (WT) mosquitoes respectively. PCR confirmation of cassette orientation was performed after LIVAK DNA extraction (32) of pupal exoskeletons using primers described in Adolfi et al., 2019 (15).

#### 2.2.1 Homozygous cassette ‘exchange’ line establishment: UAS-3A.hom

Pupal F2 progeny from UAS-3A (eYFP) and Ubi-GAL4 (CFP) crosses were screened to isolate transheterozygous individuals with eYFP/eCFP fluorescence (UAS-3A-3xP3-eYFP on one allele, Ubi-GAL4-3xP3-eCFP on the other), which were then intercrossed as adults. The F3 progeny of this cross were screened for those only carrying eYFP fluorescence as these individuals will be homozygous for the UAS-3A cassette, and designated UAS-3A.hom. Homozygosity of this line was monitored through regular screening for eYFP, and absence of non-fluorescence.

#### 2.2.2 Homozygous cassette ‘integration’ line establishment: Ubi-GAL4:UAS-3A (3A+)

‘eYFP+ eCFP+’ fluorescent pupae were selected for breeding from screens of F2 progeny from a ‘eYFP+ eCFP+ × G3’ cross and maintained as a mixed stock by enrichment of fluorescent progeny in each generation (3A+.mix). A homozygous population was later established by separating individuals based on clear differences in fluorescence intensity. Homozygosity was confirmed by out crossing to G3 and intercrossing resultant progeny without the detection of negative progeny in the next generation.

### 2.3 CCEae3A transcript expression analysis

For transcriptional and bioassay analysis, UAS-3A.hom were crossed with homozygous Ubi-GAL4 to acquire transheterozygous Ubi-GAL4/UAS-3A progeny which express CCEae3A from single alleles of the driver and responder. When required, for experiments homozygous 3A+ (3A+/3A+) were taken from stock, and heterozygous 3A+/WT and WT (WT/WT) were collected when required following YFP based screening of an unpurified mix of genotypes that were kept as a backup colony.

Three biological replicates of 3rd instar larvae and adults were collected in pools of 10 and 5 respectively in 1000μL TRIzol Reagent (Invitrogen) then RNA extracted following manufacturer’s instructions. Turbo DNA-Free kit (Ambion) and oligo(dT) SuperScript III First-Strand Synthesis System (Life Technologies) protocols were followed to remove genomic DNA from 5 μg RNA and to reverse transcribe ∼500 ng RNA aliquots respectively.

#### 2.3.1 RT-qPCR

In order to quantify the relative expression of CCEae3A between the strains generated here we undertook RT-qPCR (15). 1×Brilliant III Ultra-Fast SYBR Green qPCR Master Mix (Agilent Technologies 600882) was used with intron spanning, 3AqPCRfor (TAGCTGTCACTGTGTGGACC) and 3AqPCRrev (ACATTGTCACTGCCAGCTA) primers for assessment of CCEae3A transcript quantity. Primers qS7fw (AGAACCAGCAGACCACCATC), qS7rv (GCTGCAAACTTCGGCTATTC) (15), qEFfw (GGCAAGAGGCATAACGATCAATGCG) and qEFrv (GTCCATCTGCGACGCTCCGG) were used to quantify the housekeeping genes – ribosomal protein S7 (S7) (AGAP010592) and elongation factor (EF) (AGAP005128). 0.1 ng cDNA was included for each reaction. 3 biological and 4 technical replicates were conducted for each sample and primer pairing.

Cycle threshold (Ct) values were calculated for all samples at a baseline corrected fluorescence (dR) of 6311 up to a maximum of 35 cycles. The mean Ct (dR) was calculated for technical replicates for all samples and primer sets. Then, the mean Ct (dR) of the house keeping genes (S7 and EF) was calculated for each biological replicate. The delta Ct (ΔCt) method was used to adjust CCEae3A mean Ct values with the calculated mean Ct of the housekeeping genes for each replicate that produced a Ct for all 3 genes. The 2^-ΔΔCt method was then used to compare the expression levels between strains (calculating the fold change expression of the GOI relative to that of a second strain using normalised Ct values which had been adjusted using two housekeeping genes) and a two-tailed t-Test used to assess the significance of the difference. In control samples: Ubi-GAL4/Ubi-GAL4, Ubi-GAL4 /WT and WT/WT, both reference genes amplified at levels similar to other samples, but as expected did not amplify CCEae3A and so a ΔCt could not be calculated. Therefore, mean Ct values (with standard deviation) are reported and two tailed t-tests were used to compare homozygous and heterozygous transgenic samples values.

### 2.4 Insecticide resistance characterisation

#### 2.4.1 Larval Susceptibility Assessment

The appropriate volume of temephos, chlorpyriphos or fenthion 1×10^−4^ M stock (dissolved in acetone) to achieve the desired concentrations (2.14 nM – 2.14 μM, 24.4 nM – 1 μM, 1.37 nM - 2×10 μM respectively) was added to 200 ml of 0.01 % pondsalt water with 25 third-instar *An. gambiae* larvae in a 250 mL clear deli pot (Cater for You Ltd. SP8OZ). Mortality was assessed visually after 24 h continuous exposure. Moribund larvae were recorded as dead. 2-parameter log-logistic models (Equation 1) were generated using the ‘drc package’ (33) in R (version 4.1.0) and the comparm() function used to calculate and assess the significance of the differences in LC50.

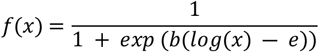

**Equation 1. Two-parameter log-logistic model**

Where the lower asymptote = 0, upper asymptote = 1, slope = b, ED50 = e

#### 2.4.2 Adult Susceptibility Assessment

WHO tube assays were used initially to test adult susceptibility using standard procedures (34). Adults were exposed for 60 min to standard diagnostic impregnated papers (Vector Control Research Unit, Universiti Sains Malaysia) of the following insecticides: OPs (malathion and pirimiphos methyl), carbamates (bendiocarb and propoxur), pyrethroids (alphacypermethrin, permethrin, deltamethrin) and OCs (dieldrin). The exposure time for DDT was adjusted to 4 h as lower exposure times did not kill most of the control adults and for fenitrothion as a 2 h exposure is recommended.

Malathion, bendiocarb and alpha-cypermethrin sensitivity was investigated further using tarsal exposure assays (35). Breifly, the compounds were dissolved in acetone and tested at concentrations from 3.9×10^−4^ – 0.3 %, 2.56×10^−6^ – 1 % and 2.56×10^−7^ – 0.1 % respectively and 0.5 mL of each was used to coat Ø60 mm Borosilicate Glass Petri Dishes (Fisher Scientific - 12901408). A 25 mL deli pot (Cater4You) with a small hole (used to aspirate mosquitoes in and out) fitting tightly into the plate creating a chamber. For each plate, 7-15x 2-5 day old female adults were aspirated through the hole, which was then covered with parafilm and mosquitoes exposed for 30min. The small size of the chamber forces tarsal contact with the insecticide. After exposure, mosquitoes were returned to cups and provided with sugar *ad libatum*, and mortality recorded 24 h later. LC50 estimates and significance were calculated using the the ‘drc’ package (33) described above.

### 2.5 Fitness cost assessment: Longevity

Whilst maintaining the 3A+/3A+ strain we noticed that there appeared to be a fitness cost in adult survival compared to other similar strains. To quantitate this apparent impact of CCEae3A expression on adult longevity, we monitored 5 replicates for each sex for 3 different generations of 3A+/3A+, compared to control Ubi-GAL4/Ubi-GAL4 *An. gambiae* strains. For each replicate, 9-11 adults were aspirated into a 200 ml paper cup and maintained with 10 % sucrose *ad libitum*. Mortality was counted and dead individuals removed every 24 h until all individuals had died. Mortality was defined as an inability to stand or fly. Differences in longevity were assessed using Kaplan-Meier Curves and Fisher exact test.

## 3 Results

### 3.1 Creation Of CCEae3A expressing *An. gambiae* by ΦC31-Mediated Transformation

CCEae3A carrying *An. gambiae* lines were generated in docking line Ubi-GAL4 (denoted A10 when originally published) (36) using site-directed ϕC31 integration of transgenes from the UAS-CCEae3A responder plasmid as described in the methods.

Supp Table 1 summarizes the embryo injection experiments that resulted in successful transformation events identified as stable fluoresence in G1 progeny derived from pooled G0 individuals. From 78 adult G0s that displayed transient larval expression of the eYFP marker, 12 G1 individuals were identified expressing one or both markers (eight eYFP – UAS-3A, four eYFP and eCFP – 3A+).

As described above, those tagged with YFP only were likely created by exchange of the Ubi-GAL4 cassette with the UAS-3A cassette to produce a regular UAS responder line. Whereas doubly tagged lines (3A+) are most likely generated by integration of the UAS-3A cassette adjacent to the Ubi-GAL4 cassette. In the 3A+ lines, ubiquitous GAL4 directed CCEae3A expression was likely to be constitutive without the need for crossing to other GAL4 driver lines.

From these 12 F1 individuals, a single isofemale UAS-3A line was selected and confirmed by PCR analysis to have a successful exchange event and was then made homozygous using the crossing and selection scheme outlined in the methods. This line was crossed with the Ubi-GAL4 line to examine CCEae3A transcript levels and insecticide resistance phenotypes described below.

One 3A+ isofemale line, confirmed by PCR to have a successful integration event was made homozygous (3A+/3A+) by intercrossing and selections based on intensity of eYFP expression. This line was analysed for CCEae3A transcript levels and insecticide resistance phenotypes without crossing to driver lines.

### 3.2 CCEae3A is Highly Transcribed in Transgenic Larvae and Adults

CCEae3A transcription in the transgenic lines was compared relative to the highly expressed housekeeping gene standards, ribosomal protein S7 (AGAP10592) and elongation factor 1 (AGAP005128). As can be observed in Fig 2, the CCEae3A is transcribed at levels similar to or higher than that of the housekeeping genes in 3rd instar larvae and adult stages in both heterozygous and homozygous conditions. Transcription is greatest in the homozygous 3A+/3A+ line, which is 8-14 fold higher than in heterozygous 3A+/WT or Ubi-GAL4/UAS-3A transheterozygote larvae (p=0.0045). Whereas, in adults the difference in transcription observed between individuals harbouring one copy and two copies of driver and responder was 4-8 fold (p=0.058 and p=0.087 for Ubi-GAL4/UAS-3A and 3A+/WT respectively). No significant difference was observed between the alternative heterozygous strains at either larvae or adult stage (p=0.199 and 0.204 respectively).

**Fig 2.**
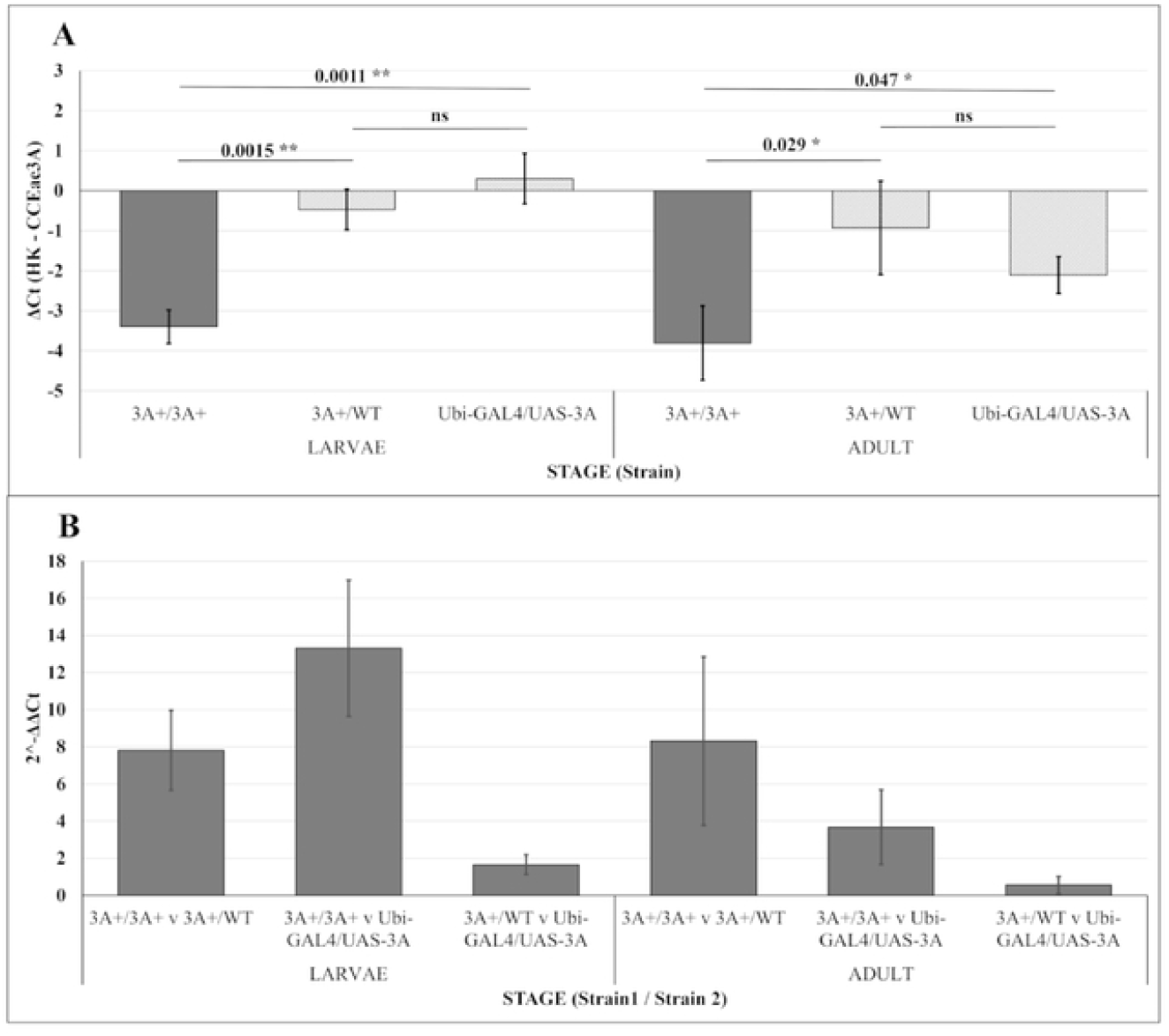
qPCR results for the CCEae3A GAIA-UAS strains confirming expression of CCEae3A In the genetically modified strains. (A) mean ΔCt values (mean Ct Housekeeping genes - Ct CCEae3A) of replicates for each stage and strain pairing. (B) mean 2^-(ΔΔCt) – comparing the CCEae3A expression between different strains which are noted below each bar separated by “v”. Values above bars indicate the p-value from a two-tailed t-test (p-value: 0.001 < ** < 0.01 < * < 0.05 < ns). Error bars on all plots = ± standard deviation of the mean.

### 3.3 Larval temephos resistance correlates with transcription level of CCEae3A

To examine potential temephos resistance conferred by CCEae3A expression, WHO larval dose response assays were initially performed with the alternative lines. The homozygous 3A+/3A+ line had an LC50 (temephos) of 198 nM, compared to 44.7 nM and 50.1 nM for heterozygous 3A+ and Ubi-GAL4/UAS-3A transheterozygote larvae respectively (Fig 3). Susceptible controls, with either Ubi-GAL4 or wildtype (WT) backgrounds, had LC50s of 35.7 nM and 33.1 nM. This translates as resistance ratios (RR) of 6.0 (p = 2.71×10-6) and 4.4 (p = 7.53×10-6) for the homozygous 3A+/3A+ against WT and heterozygous 3A+/WT respectively. The heterozygous 3A+/WT and Ubi-GAL4/UAS-3A had RRs of 1.35 (p = 0.048) and 1.40 (p = 0.068) against their respective susceptible controls (Fig 3).

**Fig 3.**
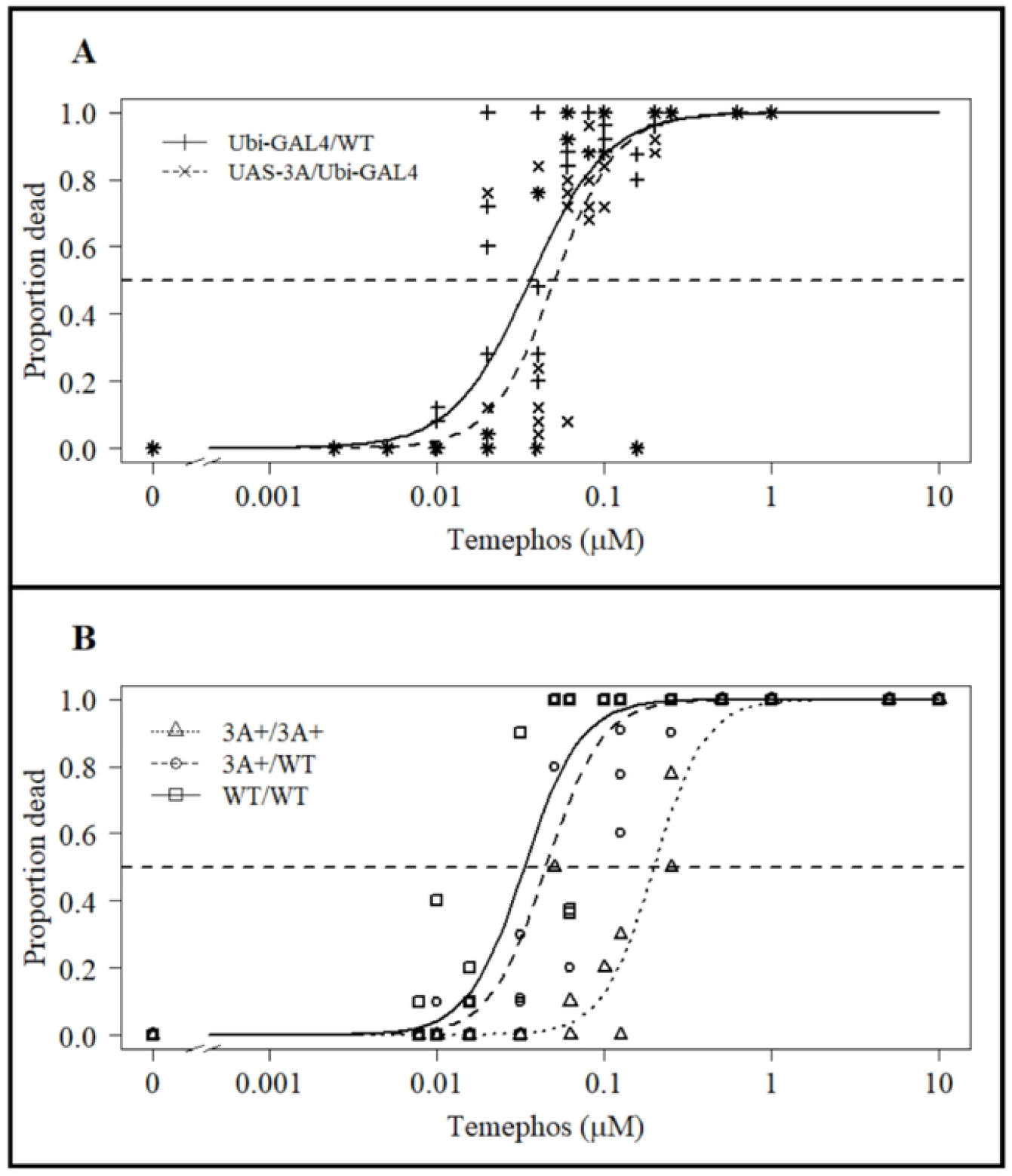
Temephos WHO larval assay results testing 3rd instar larval susceptibility for strains expressing different levels of CCEae3A. (A) Larval temephos susceptibility of the Ubi-GAL4/UAS-3A bipartite system cross (n=5) and Ubi GAL4/WT (n = 5). (B) Larval temephos susceptibility of the different genotypes of the “cassette integration” line - 3A+/3A+ (n=2), 3A+/WT (n=3) and WT/WT (n=3). Horizontal dashed line indicates y value (0.5) used for calculation of LC50s. Points are mean values for each replicate of the tested concentrations.

Since the heterozygous lines only gave low RRs, further insecticide assays were performed solely with the homozygous 3A+/3A+ line.

### 3.4 CCEae3A overexpressing larvae show resistance to other common organophosphate larvicides

In WHO dose response assays with chlorpyriphos (Fig 4A), homozygous 3A+/3A+ larvae displayed a significant (p = 5.64×10^−5^) RR of 6.6 compared with susceptible controls (LC50s of 138 nM and 20.7 nM respectively). In similar assays with fenthion (Fig 4B), homozygous 3A+/3A+ larvae showed a significant (p = 6.1×10^−12^) RR of 3.2 compared with susceptible controls (LC50s of 371 nM and 117 nM respectively).

**Fig 4.**
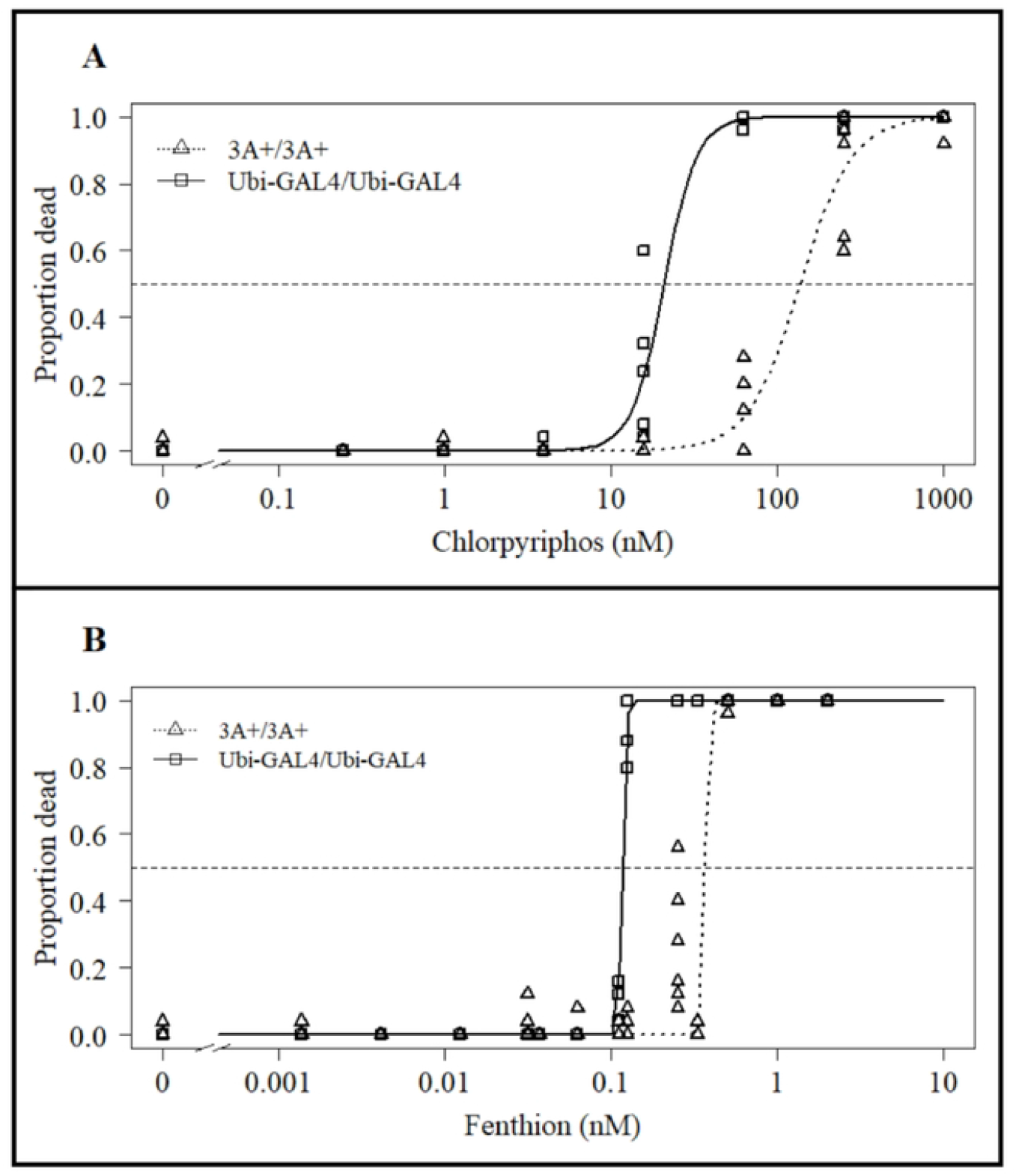
Chlorpyriphos and fenthion WHO larval assay results. Comparing 3A+/3A+ with Ubi-GAL4/Ubi-GAL4 larval susceptibility to (A) chlorpyriphos and (B) fenthion. Horizontal dashed line indicates y value (0.5) used for calculation of LC50s. Points are mean values for tested concentrations for chlorpyriphos (n = 6) and fenthion (n = 9).

### 3.5 CCEae3A expressing homozygous 3A+ adults show resistance to organophosphates, carbamates and one pyrethroid

Resistance in CCEae3A expressing adults was assessed by screening against a panel of public health insecticides using standard WHO tube diagnostic bioassays. As observed in Fig 5, significant levels of resistance (<7 % mortality) was observed against all organophosphates (fenitrothion, pirimiphos methyl and malathion) and carbamates (propoxur and bendiocarb) at standard diagnostic concentrations. There were no significant differences between CCEae3A expressing adults and susceptible controls when screening organochlorines, DDT and dieldrin, and the pyrethroids, permethrin and deltamethrin, with close to 100 % susceptibility in all assays. An inconclusive result was obtained with alphacypermethrin, which gave 90 % mortality in the CCEae3A expressing adults Fig 5, and so it was included in dose response analyses to determine accurate resistance ratios conferred by carboxylesterase expression.

**Fig 5.**
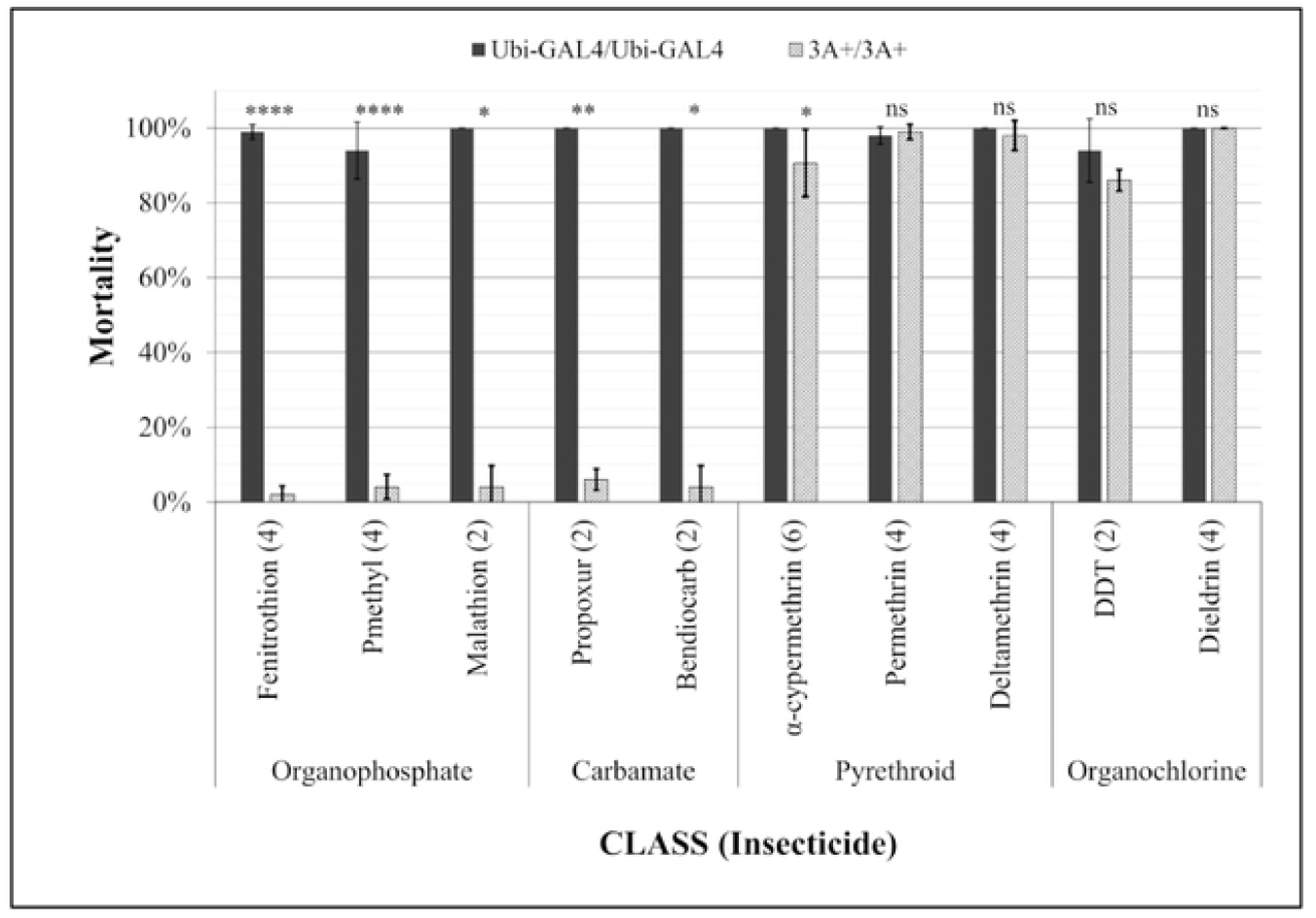
CCEae3A overexpression leads to resistance against OP and carbamates but not pyrethrolds or OCs. Adults were exposed to all compounds for I h except fenitrothion for which the standard exposure time is 2 h and DDT for which a 4 h exposure was required to kill most control mosquitoes. Error bars = standard deviation. Welch’s T-test results comparing 3A+/3A+ to Ubi-GAL4/Ubi-GAL4 for each compound are indicated above the respective bars using symbols as follows: p value - **** < 0.0001, *** < 0.001, **< 0.01, *< 0.05, · < 0.1, ns > 0.1. Numbers in brackets after compound name indicates the number of tubes tested for each strain for that compound (tested in tubes of ∼25 females).

Using a tarsal contact assay (35), adult females were exposed to a range of concentrations of test insecticides dried onto glass plates for 30 minutes and mosquito mortality scored 24 h later (Fig 6). LC50 estimates for the CCEae3A expressing adults (3A+/3A+) were 0.04 % (malathion – Fig 6A), 7.0×10^−4^ % (bendiocarb – Fig 6B), and 4.5 × 10^−4^ % (alphacypermethrin – Fig 6C), compared with 0.001 %, 3.8×10^−5^ % and 4.6×10^−5^ % respectively for the control mosquitoes. These gave corresponding resistance ratios of 36 (p < 2.2×10^−16^), 19 (p = 8.9×10^−6^) and 10 (p = 1.5×10^−5^) for the three insecticides respectively.

**Fig 6.**
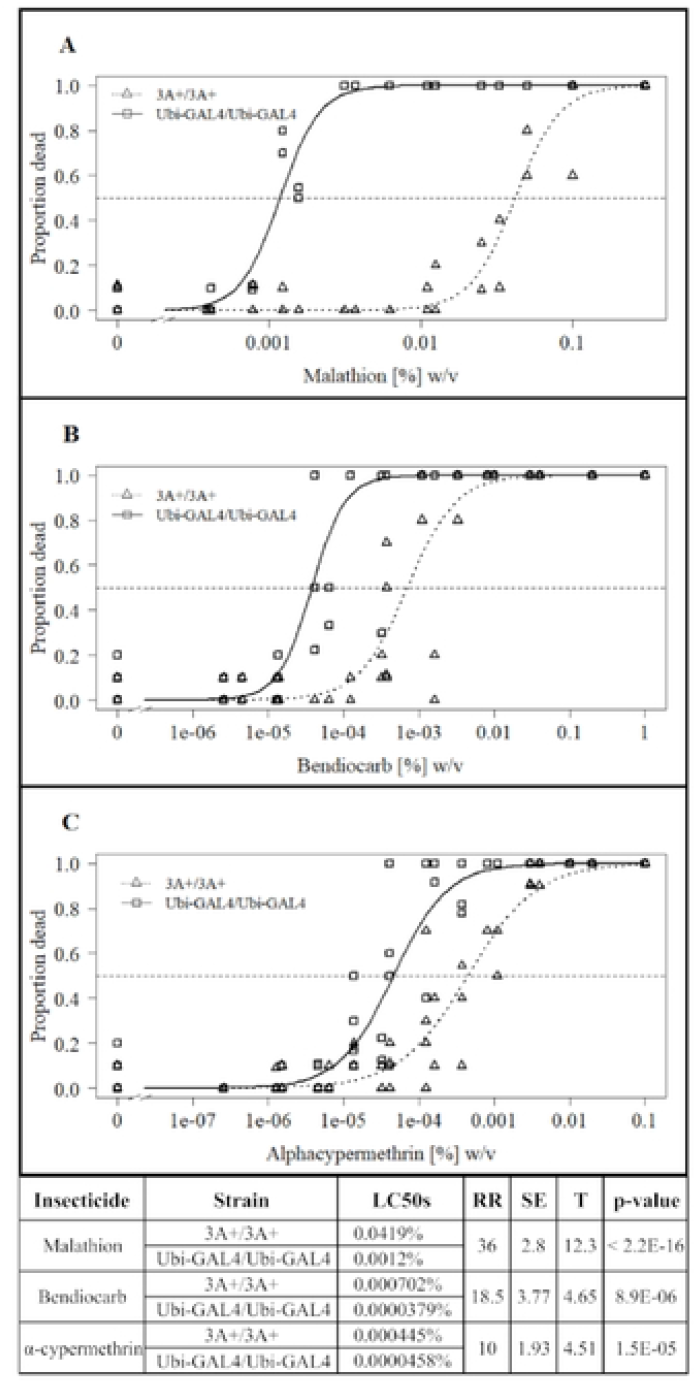
Impact of CCEae3A expression on malathion, bendiocarb and alpha-cypermethrln tarsal assay results. (A) Malathion (n=4), (B) bendiocarb (n =6) and (C) alphacypermethrin (n=6) adult tarsal assay. Lines indicate LL.2 model fit plots comparing 3A+/3A+ with Ubi-GAL4/Ubi-GAL4. Horizontal dashed line indicates y value (0.5) used for calculation of LC50s. Points are mean values for each replicate of the tested concentrations. (D)Table detailing statistical outcomes of Z-test analysis comparing the LC50 values of 3A+/3A+ and Ubi-GAL4/Ubi-GAL4.

### 3.6 CCEae3A expressing homozygous 3A+/3A+ adults show reduced longevity

To determine if there were fitness costs associated with expressing CCEae3A, survival assays were performed to measure adult longevity of both sexes compared to parental control mosquitoes. Log rank analysis indicated significant impact of both strain (p = 0.00018) and sex (p = 0.00025). Further log rank analysis was run to assess the difference between homozygous CCEae3A and control mosquitoes for each of the sexes separately. Significant differences were found for both females (Fig 7A, p = 0.00021) and males (Fig 7B, p = 0.035), although median survival time did not vary for females and was only one day different for males. CCEae3A females appear to survive at similar rates to controls until day 17, then have an increased death rate thereafter. Conversely, CCEae3A males die off more rapidly until 17 days, but thereafter have similar rates of death.

**Fig 7.**
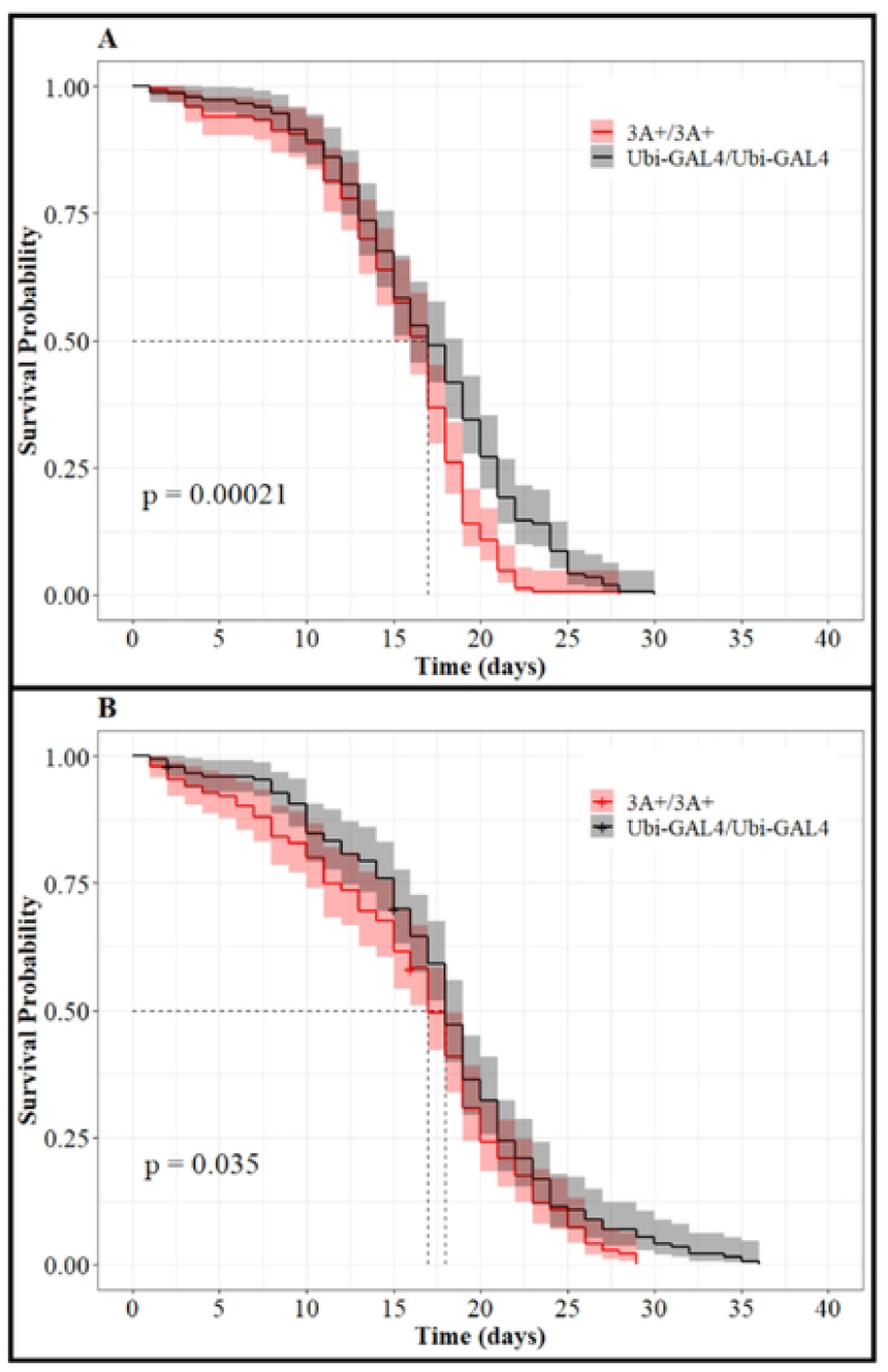
Impact of CCeae3A expression on longevity. Kaplan-Meier plots of 3A+/3A+ and Ubi-GAL4/Ubi-GAL4 for (A) female and (B) males displaying the impact of CCEae3A expression and sex on longevity. P-values reported are the result of a log rank test comparing the curves in each panel. Shadows represent the 95 % confidence intervals for each day. Dotted line indicates the mean survival probability for each strain.

## 4 Discussion

CCEae3A overexpression and duplication has been reported in multiple insecticide cross resistant *Ae. aegypti* populations across the world (22-25, 27-29). In studies of these recently colonised or lab selected mosquito strains it is difficult to quantify the role of individual genes in conferring a specific resistance phenotype because multiple resistance mechanisms and multiple insecticide resistance phenotypes co-exist. Here, we have shown that transgenic expression of the *Ae. aegypti* carboxylase CCEae3A in an otherwise susceptible *An. gambiae* genetic background is sufficient to induce high levels of cross resistance (3-36 fold) to a range of ester bond containing insecticides comprising of OPs, including the widely used larvicide temephos, and the carbamate adulticides, bendiocarb and propoxur. Moreover, pyrethroid resistance was also observed in CCEae3A expressing transgenic lines but was restricted to alphacypermethrin, and not to permethrin and deltamethrin.

It must be borne in mind though that the experiments were performed in *An. gambiae*, and we can’t rule out that fine differences in resistance phenotypes may have been observed if performed in Aedes. We chose to use the GAL4/UAS system developed in *An. gambiae*, since setting up a similar system in Aedes from scratch would be very time consuming. Moreover, we have a panel of *An. gambiae* responder lines available and so can directly compare the levels and range of resistance provided by carboxylase expression with those of P450s, GSTs and Sap2 in our panel (14,15). In contrast to the other panel members, the CCEae3A line provides high level resistance to both carbamates and OPs. This considerably broadens the range of metabolic activity of the panel for use in screening the liability of novel compounds coming to market (17).

In adults, although not statistically significant, 4-8 fold greater transcripts were observed in the homozygous line which carries 2 copies of both pUb-GAL4 and UAS-CCEae3a when compared with the each of the heterozygous lines which possess a single copy of each. Larvae with two copies of the GAL4 and UAS 3A transgenes gave 8-14 fold higher transcription yield of CCEae3A than those with one copy. In such a binary system as GAL4/UAS, one would expect a two-four fold increase (dependant on the saturation of the components) however since we see higher levels, it may suggest the occurrence of an activating interaction (transvection) between the transgene on one chromosome and the corresponding transgene on the homologous chromosome (37).

Initial experiments with temephos indicated that the magnitude of resistance in the CCEae3A lines was correlated with gene dosage and consequent transcription level. The homozygous Ubi:3A displayed an increase in larval temephos resistance 4-6 fold compared to the heterozygous counterparts. It was noted that the level of resistance in dual copy strains was only marginally ∼1.5 fold greater than susceptible controls. Which would suggest that a threshold of CCEae3A expression may be required to confer significant resistance to temephos by WHO standards.

The temephos resistance observed in the transgenic lines provides further support to *in vitro* studies which show that CCEae3A is capable of sequestering and metabolising the temephos-oxon (26) and that upregulation of CCEae3A correlates with temephos resistance in the NK2 strain from which the transgene was cloned (22). The RR for NK2 and 3A+/3A+ against their respective susceptible controls were of similar magnitude, NK2 6-10-fold and 3A+/3A+ 6-fold. In other tempehos resistant strains associated with CCEae3A upregulation, RRs of 2.3-fold (25), 13-36-fold (38) and 15-29-fold (23) have been reported, perhaps indicating the presence of other resistance factors in these highly resistant wild strains.

Larvae of the homozygous Ubi:3A strain also displayed resistance to both chlorpyriphos (6.64-fold) and fenthion (3.18-fold) OPs. As far as we are aware, the larval expression of CCEae3A has not been directly linked to chlorpyriphos and fenthion resistance. Although the role of esterases has been indicated by synergist studies in a chlorpyriphos resistant *Ae. aegypti* line (39), and raised esterase activity in fenthion resistant Culex mosquitoes (40).

This broad spectrum activity against OPs was also shown in adult bioassays. CCEae3A overexpression gave rise to high resistance to malathion (RR = 36), fenitrothion (<5 % WHO mortality) and pirimiphos-methyl (<5 % WHO mortality). Only recently has pirimiphos methyl gained registration for use in household vector control programmes for IRS (41) so these findings are very concerning. Monitoring of the already documented resistance to pirimithos methyl conferred through insensitive acetylcholine esterase in new exposed vector populations (42) should also be aware of the potential co-evolution of carboxylesterases to synergise the resistance profile.

Meanwhile, the other OP compounds have been used extensively for adult control. In wild caught populations, CCEae3A upregulation (1.8-11.1 fold) correlated with low levels (70-90% WHO mortality (29)) or no (27) fenitrothion resistance in Senegal and Madeira Island respectively. This suggest that higher levels of CCEae3A upregulation may be necessary to cause resistance to other OPs than that which is required to provide substantial malathion resistance.

Malathion resistance has been linked several times previously to CCeae3A in field caught mosquitoes (23, 25, 29, 43) and also to other insect carboxylesterase enzymes, including transgenic *D. melanogaster*, in which LcαE7 Rmal mutant (44) and CpEST (wildtype) (45) overexpressing flies displayed ∼30 and 1.5 fold malathion resistance respectively. The combination of malathion and temephos resistance conferred by CCEae3A is concerning as although many countries are adopting rotational or mosaic combinations of different insecticides for insecticide resistance management (46), these insecticides are often still crucial components due to the lack of alternative effective and approved compounds.

Homozygous 3A+/3A+ mosquitoes also displayed extremely high resistance to both carbamates tested, propoxur (<7 % mortality) and bendiocarb (RR = 19). Our data support the role of CCEae3A upregulation causing the WHO diagnostic resistance to bendiocarb (< 63% mortality) and propoxur (<78 % mortality) which was observed in *Ae. aegypti* collected in Senegal (29). Bendiocarb resistance (60-75% mortality) has also been observed in the Maderia Island *Ae. aegypti* with upregulated CCEae3A (27). In these mosquitoes, nearly 100% mortality was observed when co-exposed with PBO, and it was concluded that the resistance was largely the due to P450 metabolism. As little resistance to fenitrothion was also observed in the Madeira population, the limited CCEae3A upregulation observed was likely not impacting greatly on the carbamate resistance profile in these mosquitoes. In WHO assays, homozygous 3A+/3A+ displayed full susceptibility to permethrin and deltamethrin despite both compounds containing ester groups. There has been no evidence in previous studies of CCEae3A upregulation that directly contradicts these results (23, 25, 27, 29, 38). In these previous studies, pyrethroid resistance in mosquitoes overexpressing CCEae3A was more strongly associated with other classes of known pyrethroid metabolising enzymes, particularly P450s, which were also overexpressed.

Potential resistance to alphacypermethrin (91 % mortality) was detected in diagnostic WHO assays, during which it was observed that 3A+/3A+ resisted knockdown for longer times than controls. A RR of 9.71 was then confirmed though dose response assays. This pyrethroid phenotype has not been associated with CCEae3A previously. This data would suggest that the permethrin resistance seen in various *Ae. aegypti* populations from Martinique, Madeira, and Guadeloupe is not linked to CCEae3A up regulation in these cross resistant populations. Whereas the alphacypermethrin resistance observed in *Ae. aegypti* populations from Senegal (29) may well be influenced by the reported CCEae3A overexpression. Alphacypermethrin, which is primarily composed of the most active cis isomers of cypermethrin, contains an ester group which has been shown to be cleaved during toxicity studies in mammals, presumably by carboxylesterases (47). Alpha-esterases have been implicated in pyrethroid resistance previously in *Ae. aegypti* (48, 49) though even in these cases the role of alpha-esterases has been questioned. In previous mosquito studies, when CCEae3A upregulation and alphacypermethrin resistance have been co-detected (29) other mechanisms of resistance have also been present and the alphacypermethrin resistance has been attributed entirely to mutations in the voltage gated sodium channel encoding gene (*kdr* mutants) and cytochrome P450 based resistance mechanisms (50).

Most research into combatting pyrethroid resistance has been conducted on Anopheles mosquitoes, however PBO resistance has been reported previously in *Ae. aegypti* in Florida Keys (51) and although resistance was not detected to permethrin and deltamethrin herein, the alphacypermethrin resistance detected here is concerning, since it is also a widely used pyrethroid. The potential role of carboxylesterases should be borne in mind when monitoring the effectiveness of pyrethroid resistance breaking compounds that circumvent P450 upregulation such as piperonyl butoxide (PBO) (52) inclusion on bed nets and when alphacypermethrin is used for Ae aegypti control to combat resistance to OPs.

As expected, no resistance to organochlorines, DDT or dieldrin, was detected in the CCEae3A upregulated mosquitoes, as these molecules lack a carboxylester target group.

During maintenance of the constitutively expressing CCEae3A line, anecdotal observation indicated that there was a longevity fitness cost in that Ubi:3A mosquitoes did not seem to survive as long as other lines kept in the insectary. To quantitate this, accurate mortality counting was performed which indicated that although there were only minor differences in median survival to controls, significant reduction in survival rates over time did occur. 3A+/3A+ female death increases rapidly after day 17 resulting in fewer numbers compared to controls after 22 days. Whereas in males, there was a bimodal pattern of death. In the first week, 3A+/3A+ preferentially died, but survived better thereafter until day 25, when relative die off increased again. Since we examined insecticide resistance in 3-7 day old females, this mortality is unlikely to have affected the resistance profiling. The impact of CCeae3A upregulation on longevity has not been specifically investigated previously but there have been reports of reduced longevity in temephos resistant female *Ae. aegypti* from Brazil that displayed increased activity of esterases (53). However, it is unclear how the carboxylase or GAL4 expression exerts its fitness cost in a sex specific manner and would be interesting to follow up.

By generating both the cassette integration line (3A+/3A+) and the cassette exchange line (UAS-3A.hom) we can combine CCEae3A overexpression with overexpression of other genes or resistance mechanisms such as kdr mutations and of testing a variety of different promotors to vary the localisation and level of expression using the two different lines respectively. This allows us to investigate in detail the effects of CCEae3A overexpression alone and to study potential synergism between different mechanisms which was found to be important in the past in mosquitoes with kdr mutation which also overexpressed GSTe2. These mosquitoes were found to be more resistant to permethrin than those carrying kdr which did not overexpress GSTe2 although GSTe2 overexpressing mosquitoes without the kdr mutation were fully susceptible to permethrin (17).

The role of CCEae3A overexpression in resistance to several of the compounds tested was unclear prior to this study as insecticide selection results in multiple molecular changes which can be difficult to unravel. Here we demonstrate the importance of investigating and characterising suspected resistance mechanisms in isolation to accurately characterise the potential effect. This will become increasingly important as around the world countries adopt resistance management practices such as insecticide rotation and molecular screening of resistant populations for ‘known’ resistance markers. Poor understanding of cross resistance and of the role of individual molecular mechanisms could result in failures to curb resistance spread and reduced efficacy of mosquito control programmes.

## Acknowledgements

We thank Dr Frank Meechan and Dr James Maas for their assistance with data analysis and Professor Hilary Ranson for supply of the original Nakhon Sawan 2 *Ae. aegypti* CCEae3A cDNA plasmid. The work was funded through the MRC CASE studentship MR/P016197/1 awarded to GJL and DBS.

